# Intracellular DNA replication and differentiation of *Trypanosoma cruzi* is asynchronous within individual host cells *in vivo* at all stages of infection

**DOI:** 10.1101/2019.12.20.884221

**Authors:** Martin. C. Taylor, Alexander Ward, Francisco Olmo, Shiromani Jayawardhana, Amanda F. Francisco, Michael D. Lewis, John M. Kelly

## Abstract

Investigations into intracellular replication and differentiation of *Trypanosoma cruzi* within the mammalian host have been restricted by limitations in our ability to detect parasitized cells throughout the course of infection. We have overcome this problem by generating genetically modified parasites that express a bioluminescent/fluorescent fusion protein. By combining *in vivo* imaging and confocal microscopy, this has enabled us to routinely visualise murine infections at the level of individual host cells. These studies reveal that intracellular parasite replication is an asynchronous process, irrespective of tissue location or disease stage. Furthermore, using TUNEL assays and EdU labelling, we demonstrate that within individual infected cells, replication of both mitochondrial (kDNA) and nuclear genomes is not co-ordinated within the parasite population, and that replicating amastigotes and non-replicating trypomastigotes can co-exist in the same cell. Finally, we report the presence of distinct non-canonical morphological forms of *T. cruzi* in the mammalian host. These appear to represent transitional forms in the amastigote to trypomastigote differentiation process. Therefore, the intracellular life-cycle of *T. cruzi in vivo* is more complex than previously realised, with potential implications for our understanding of disease pathogenesis, immune evasion and drug development. Dissecting the mechanisms involved will be an important experimental challenge.

**AUTHOR SUMMARY:** Chagas disease, caused by the protozoan parasite T*rypanosoma cruzi*, is becoming an emerging threat in non-endemic countries and establishing new foci in endemic countries. The treatment available has not changed significantly in over 40 years. Therefore, there is an urgent need for a greater understanding of parasite biology and disease pathogenesis to identify new therapeutic targets and to maximise the efficient use of existing drugs. We have used genetically modified strains of *T. cruzi* carrying a bioluminescence/fluorescence dual reporter fusion gene to monitor parasite replication *in vivo* during both acute and chronic infections in a mouse model. Utilising TUNEL assays for mitochondrial DNA replication and EdU incorporation for total DNA replication, we have found that parasite division within infected cells is asynchronous in all phases of infection. Differentiation also appears to be uncoordinated, with replicating amastigotes co-existing with non-dividing trypomastigotes in the same host cell.

## INTRODUCTION

The obligate intracellular parasite *Trypanosoma cruzi* is responsible for Chagas disease, a debilitating infection that is widespread in Latin America. There are an estimated 6-7 million people infected [1]. In addition, due to migration, cases are increasingly being detected outside endemic regions [2, 3]. *T. cruzi* is spread by blood-sucking triatomine bugs, although oral transmission via contaminated food or drink, and the congenital route are also important. The parasite has a wide mammalian host range and can infect most nucleated cells. During its life-cycle, the major features of which were established more than a century ago [4], *T. cruzi* passes through a number of differentiation stages involving both replicative and non-replicative forms. Infections are initiated by insect transmitted metacyclic trypomastigotes, which are flagellated and non-replicating. Once these have invaded host cells, they escape from the parasitophorous vacuole into the cytosol, differentiate into ovoid non-motile amastigotes, and divide by binary fission. After a period of approximately 4-7 days, by which time parasite numbers can have reached several hundred per infected cell, they differentiate into non-replicating flagellated motile trypomastigotes. This eventually promotes host cell lysis, and the released parasites then invade other cells, spread systemically through blood and tissue fluids, or can be taken up by triatomine bugs during a bloodmeal. Within the insect vector, they differentiate into replicating epimastigotes, and finally metacyclic trypomastigotes, to complete the cycle.

More recently, *in vitro* studies have suggested that the parasite life-cycle may be more complex than outlined above. These reports include the identification of an intracellular epimastigote-like form [5], and amastigote-like forms with short flagella, termed sphaeromastigotes [6]. Whether these parasite forms represent intermediate transitional types, or correspond to intracellular stages with a specific role, remains to be determined. Adding to the complexity, trypomastigotes can also differentiate into an epimastigote-like stage, via an amastigote-like transitional form [7]. These recently differentiated epimastigotes have a distinct proteomic profile, display complement-resistance, can invade phagocytic and cardiac cells, and are infectious to mice. In addition, it has been reported that when bloodstream trypomastigotes invade mammalian cells, they can undergo a differentiation step in which asymmetric cell division results in the generation of an amastigote, together with a second, defective parasite cell termed a zoid, which contains a kinetoplast, but lacks a nucleus [8]. This has not, as yet, been demonstrated for the metacyclic trypomastigote which initiates natural mammalian infection. Most recently, it has been observed that infrequent spontaneous dormancy can occur in intracellular amastigotes, a phenomenon that may be linked to increased drug tolerance [9]. These non-proliferating intracellular amastigotes, which have been identified both *in vivo* and *in vitro*, retain the ability to differentiate into trypomastigotes. Their metabolic status is unknown. To date, a lack of sufficiently sensitive *in vivo* parasite detection methods has meant that it has not been possible to investigate the biological role of these and the other non-classical parasite forms during either acute or chronic stage infections.

There are three distinct stages to Chagas disease. In humans, the acute stage occurs in the first 4-6 weeks, and is characterised by a widely disseminated infection, together with patent parasitemia. This results in the induction of a robust CD8^+^ T cell-mediated response [10], with infected individuals then progressing to the asymptomatic chronic stage, where the parasite burden is extremely low and difficult to detect. Around 30-40% of those infected eventually develop chronic disease pathology, predominantly cardiomyopathy and/or digestive tract megasyndromes [11, 12]. In humans, infections with *T. cruzi* are considered to be life-long, however our understanding of parasite biology and tropism during the chronic stage, and their relationship to disease outcome are limited [13]. To address these issues, we developed an experimental murine model based on highly sensitive bioluminescence imaging of *T. cruzi* genetically modified to express a red-shifted luciferase [14, 15]. This system allows chronic infections to be followed in real time for periods of longer than a year, and enables endpoint assessment of parasite location by *ex vivo* imaging. In this mouse model, the infection is pan-tropic during the acute stage and parasites are readily detectable in almost all organs and tissues. During the chronic stage however, the parasite burden is very low and restricted mainly to the colon and/or stomach, with other organs/tissues infected only sporadically [14, 16].

Although bioluminescence can be widely used for *in vivo* testing of drugs and vaccines, and as a technique for exploring infection kinetics and dynamics, it does not easily allow the identification or study of single parasites at a cellular level [16-19]. To overcome this limitation, we re-engineered the *T. cruzi* strain to express a bioluminescent/fluorescent fusion protein [20]. The aim was to enable infection dynamics to be monitored at a whole animal level using bioluminescence, followed by investigation of host-parasite interactions at a single cell level using fluorescence. With this approach, we have been able to routinely image individual parasites in murine tissues during chronic stage infections. This has allowed us to readily visualise parasites residing within individual host cells in chronically infected animals. Here, we describe the exploitation of this dual imaging procedure to gain new insights into parasite biology in experimental models of acute and chronic Chagas disease.

## METHODS

### Parasite culture

*T. cruzi* CL-Luc::Neon epimastigotes were cultured in supplemented RPMI-1640 as described previously [21]. Genetically manipulated lines were routinely maintained on their selective agent (hygromycin, 150 μg ml^-1^; puromycin, 5 μg ml^-1^; blasticidin, 10 μg ml^-1^; G418, 100 μg ml^-1^). MA-104 (fetal African green monkey kidney epithelial) cells (ATCC CRL-2378.1) were cultivated to 95–100% confluency in Minimum Essential Medium Eagle (MEM, Sigma.), supplemented with 5 % Foetal Bovine Serum (FBS), 100 U/ml of penicillin, and 100 μg ml^-1^ streptomycin at 37°C and 5% CO_2_. Tissue culture trypomastigotes (TCTs) were derived by infecting MA104 cells with stationary phase metacyclic trypomastigotes. Cell cultures were infected for 18 hours. External parasites were then removed by washing in Hank’s Balanced Salt Solution (Sigma-Aldrich), and the flasks incubated with fresh medium (Minimum Essential Medium (Sigma-Aldrich) supplemented with 5% FBS) for a further 5-7 days. Extracellular TCTs were isolated by centrifugation at 1600 *g*. Pellets were re-suspended in Dulbecco’s PBS and motile trypomastigotes counted using a haemocytometer. *In vitro* infections for microscopy were carried out as above, but on coverslips incubated in 24-well plates using an MOI of 5:1 (host cell:parasite). Coverslips were fixed with 2% paraformaldehyde at 72 hours post infection. Cells were then labelled with TUNEL (section 4.6).

### Ethics statement

All animal work was performed under UK Home Office licence 70/8207 and approved by the London School of Hygiene and Tropical Medicine Animal Welfare and Ethical Review Board. All protocols and procedures were conducted in accordance with the UK Animals (Scientific Procedures) Act 1986.

### Mouse infection and necropsy

Mice were maintained under specific pathogen-free conditions in individually ventilated cages. They experienced a 12 hour light/dark cycle and had access to food and water *ad libitum*. Female mice aged 8-12 weeks were used. CB17 SCID mice were infected with 1×10^4^ tissue culture trypomastigotes, and monitored by bioluminescence imaging (BLI), as previously reported [14]. At the peak of the bioluminescence signal, when trypomastigotes were visible in the bloodstream, the mouse was culled by an overdose of pentobarbital sodium, and the infected blood obtained by exsanguination. The trypomastigotes were washed in Dulbecco’s PBS and diluted to 5×10^3^ ml^-1^. 1×10^3^ trypomastigotes were injected i.p. into each mouse (BALB/c or C3H/HeN) and the course of infection followed by BLI. At specific time-points, the mice were euthanised by an overdose of pentobarbital sodium and necropsied (for detailed description of the necropsy method, see Taylor *et al*., 2019). Their organs were subject to *post mortem* BLI. We excised those segments that were bioluminescence-positive and placed them into histology cassettes. BLI images from living animals and *post-mortem* tissues were analysed using Living Image 4.5.4 (PerkinElmer Inc.)

### Tissue embedding and sectioning

Tissue sections were produced as described previously [20, 22]. Briefly, excised tissue was fixed in pre-chilled 95% ethanol for 20-24 hours in histology cassettes. The tissues were dehydrated in 100% ethanol, cleared in xylene, and then embedded in paraffin at 56°C. Sections were cut with a microtome and mounted on glass slides, then dried overnight. Slides were stored in the dark at room temperature until required.

### TUNEL assay for kDNA replication

For *in vitro* studies, logarithmically growing epimastigotes and infected mammalian cells on coverslips were fixed with 2% paraformaldehyde in PBS. Fixed epimastigotes were air-dried onto glass 8-well slides. Cells were washed once in PBS and permeabilized in 0.1% TritonX-100/PBS for 5 min and washed 3 times with PBS. 20 µL TUNEL reaction mixture (In situ Cell Death Detection Kit, TMR-red, Roche) was added to each well or coverslip and the reaction incubated for 1 hour at 37°C. For tissue sections, slides were deparaffinised in 2 changes (30 s each) of xylene, 3 changes (1 min each) of pre-chilled 95% ethanol, and 3 changes (1 min each) of pre-chilled Tris-buffered saline (TBS). Sections were outlined with a hydrophobic pen then permeabilized in 0.1% TritonX-100/PBS for 5 min and washed 3 times with PBS. 20 µL TUNEL reaction mixture was added to each section and the slide was overlaid with a coverslip to ensure that the reaction mix was evenly distributed. The reaction was incubated for 20 min to 2 hours at 37°C. Coverslips and slides were mounted in VECTASHIELD® with DAPI (Vector Laboratories, Inc.) before observation on a Zeiss Axioplan LSM510 confocal microscope.

### EdU assay for DNA replication

Mice were injected intraperitoneally with 12.5 mg kg^-1^ EdU (Sigma-Aldrich) in PBS at specific time points (as detailed in Results) prior to euthanasia. Tissues were fixed and sectioned as above. Labelling of the incorporated EdU was carried out using the Click-iT Plus EdU AlexaFluor 555 Imaging kit (Invitrogen), following a similar method as used for TUNEL labelling, but substituting the Click-iT reagent for the TUNEL reaction mix. For sections which had been in paraffin for extended time periods (> 6 months), the slides were immersed in 100 mM EDTA for 16 hours (on manufacturer’s recommendation), then washed extensively with TBS prior to the Click-iT reaction.

### Confocal microscopy

Slides and sections were examined using a Zeiss LSM510 Axioplan confocal laser scanning microscope. Cells containing multiple parasites were imaged in three dimensions to allow precise counting of amastigotes (using the 63x or 100x objectives with appropriate scan zoom for the particular cell/number of parasites). Phase images were obtained at lower magnification (40x) to allow orientation of the tissue section and identification of specific layers/structures. All images were acquired using Zeiss LSM510 software. Scale bars were added using the Zeiss LSM Image Browser overlay function and the images were then exported as .TIF files to generate the Figs.

### Live imaging of infected cells

Videos were acquired using an inverted Nikon Eclipse microscope. The chamber containing the specimen was moved in the x-y plane through the 580 nm LED illumination. Images were collected using a 16-bit, 1-megapixel Pike AVT (F-100B) CCD camera set in the detector plane. An Olympus LMPlanFLN 20x/0.40 objective was used to collect the exit wave leaving the specimen. Time-lapse imaging was performed by placing the chamber slide on the microscope surrounded by an environmental chamber (Solent Scientific Limited, UK) maintaining the cells and the microscope at 37°C / 5% CO_2_. Time-lapse video sequences were created using the deconvolution app in the Nikon imaging software.

## RESULTS

### Parasite kinetoplast DNA replication is not synchronised within individual infected cells

The text book view of the *T. cruzi* intracellular cycle is that invading trypomastigotes differentiate into amastigotes, which then begin to divide by binary fission within the cytoplasm of the host cell. These then differentiate into trypomastigotes and the host cell lyses releasing the trypanosomes, see for example Fig 1a in [23]. However, the degree to which amastigote division and differentiation are co-ordinated within single cells, and the potential for this to be influenced by host cell type and/or tissue-specific location are poorly understood.

**FIG 1.**
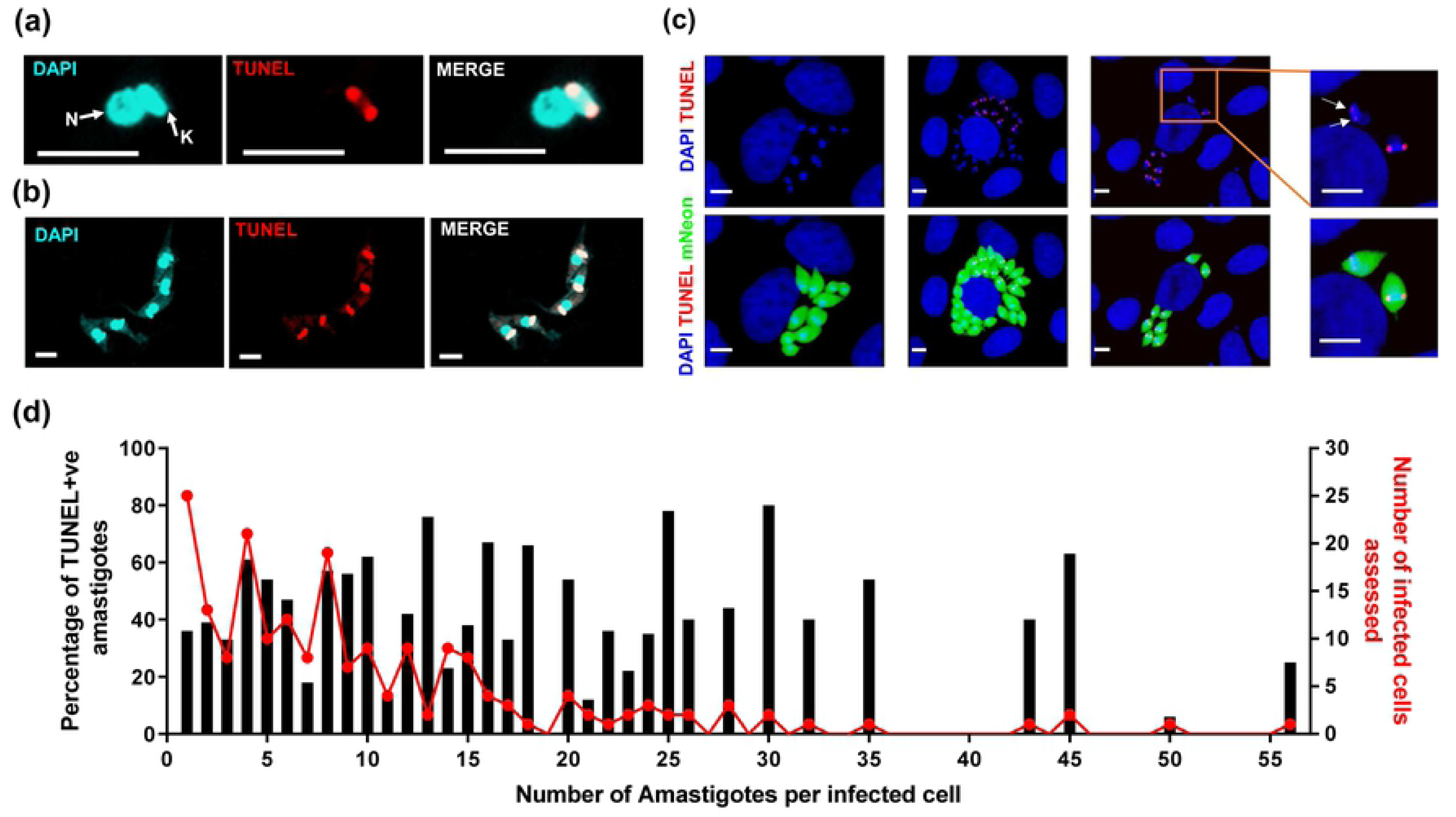
Kinetoplast replication of *T. cruzi* amastigotes is asynchronous in vitro. (a) Epimastigote at early stage of kDNA replication with TUNEL labelling of antipodal sites. (b) Epimastigotes at late stage of kDNA replication showing TUNEL labelling of entire kDNA disk. (c) MA104 cells infected with *T. cruzi* CL-Luc::Neon amastigotes for 72 hours then fixed and labelled with the TUNEL reagent. Left hand panel: cell containing 11 amastigotes with non-replicating kDNA (all TUNEL-ve); central panel: cell with parasites in which kDNA replication is asynchronous (mix of TUNEL+ve and TUNEL-ve); right hand panel: cell where all amastigotes are TUNEL+ve, but at different stages of kDNA replication (7 of 8 amastigotes display bright antipodal staining, the eighth is faintly TUNEL+ve, as shown by white arrows in the inset). (d) TUNEL data from 200 infected cells pooled from 3 replicate wells. The red line represents the number of infected cells assessed that contained the specified number of resident amastigotes. The black bars represent the percentage of amastigotes per cell that label as TUNEL+ve. Bar = 5 μm

During trypanosomatid cell division there are two distinct DNA replication events that result in duplication of the mitochondrial (kinetoplast or kDNA) and then the nuclear genomes. However, at early stages of kDNA or nuclear DNA replication, it is not feasible to assign parasites to a particular cell-cycle phase by morphology or total DNA staining, as many parasites appear similar. To identify the replication status of the mitochondrial genome in intracellular amastigotes we took advantage of the TUNEL assay (terminal deoxynucleotidyl transferase dUTP nick end labelling), a procedure normally used to quantify apoptotic cell death in mammalian cells [24]. In *T. cruzi*, this assay can be utilised to monitor kDNA replication [20], a genome that consists of thousands of catenated circular double-stranded DNA molecules. The majority of these are the mini-circles that encode the guide RNAs that mediate RNA editing [25]. To maintain functional RNA editing, daughter cells must each inherit copies of the entire mini-circle repertoire. During replication, mini-circles are first detached from the catenated network and the new strands are then synthesised. However, some of the single-strand breaks that result from removal of RNA primers in the newly synthesised DNA are maintained until the whole mini-circle network has been replicated. This enables newly duplicated circles to be distinguished from non-replicated circles and ensure each daughter network is complete [26, 27]. Therefore, during the S-phase of kDNA replication, the free 3’ hydroxyl groups at the nicks on the newly synthesised strands can be labelled with a fluorescent analogue by terminal uridylyl transferase [20, 26, 28]. This means that the TUNEL assay enables specific labelling of parasites that have commenced cell division.

We first applied TUNEL assays to asynchronous, exponentially growing epimastigote cultures to confirm that this method was applicable to *T. cruzi*. Parasites in the early phase of kDNA synthesis displayed TUNEL positivity in antipodal sites on either side of the kDNA disk, indicative of the two replication factories (Fig 1a). Later in replication, the entire disk was labelled (Fig 1b). Nuclear DNA did not exhibit a positive signal at any stage (Fig 1a and b).

To quantify the replication of kDNA in intracellular amastigotes, the parasites in 200 infected cells were assessed for TUNEL positivity *in vitro* 72 hours post-infection. These cultures were infected with a low multiplicity of infection (1 parasite per 5 host cells) to minimise the chance of individual cells being infected twice. It was apparent that kDNA replication within single infected cells was largely asynchronous, since most infected cells contained both TUNEL+ve and –ve amastigotes (Fig 1c and d). Most TUNEL+ve parasites displayed antipodal staining, indicative of early phase replication (see examples in Fig 1c). The number of amastigotes displaying whole disk staining was low suggesting that kDNA nick repair may occur more rapidly than in epimastigotes. The few amastigotes that displayed a 2K1N morphology showed no TUNEL staining on either kinetoplast, indicating that nicks are repaired prior to segregation, as expected (example shown in S1 Fig) [26].

Total amastigote numbers within infected cells were also consistent with asynchronous replication; they did not follow a geometric progression as would be expected if growth was co-ordinated (Fig 1d, red line). There were no cases where a specific number of amastigotes within a cell was always associated with 100% TUNEL labelling (S2 Fig). Intracellular populations of 2, 4 or 8 amastigotes were equally as likely to be asynchronous as populations containing non-geometric numbers (Fig 1d, black bars, S2 Fig). In the minority of infected cells where every amastigote was TUNEL+ve (14.5% of cells that contained more than one parasite), there were differences in the degree of labelling between the parasites in 24% of the host cells (Fig 1c inset, for example, white arrows indicate faint TUNEL labelling of one amastigote in earlier phase of kDNA replication). Collectively, these results therefore show that within a single infected cell *in vitro*, amastigote kDNA replication is not synchronised within the population.

We then applied the TUNEL assay to mouse tissues obtained from acute experimental infections with the dual bioluminescent/fluorescent *T. cruzi* cell line CL-Luc::Neon [20]. The acute phase in mice is characterised by widespread dissemination of infection with amastigotes in diverse cell and tissue types. We sampled a range of organs and tissues (Fig 2; S3 Fig). This revealed that within any given infected host cell, the extent of kDNA labelling varied between parasites. We quantified the frequency of TUNEL positivity amongst amastigotes in sections from various organs in a single mouse (Fig 3). The majority of amastigotes in the acute phase had TUNEL+ve kDNA, showing that they were undergoing replication. However, there was no evidence for programmed synchronicity, and in each tissue, individual cells could contain both TUNEL+ve and TUNEL-ve parasites. Moreover, all of the different organs that were analysed showed similar profiles with respect to parasite replication states (Fig 3).

**FIG 2.**
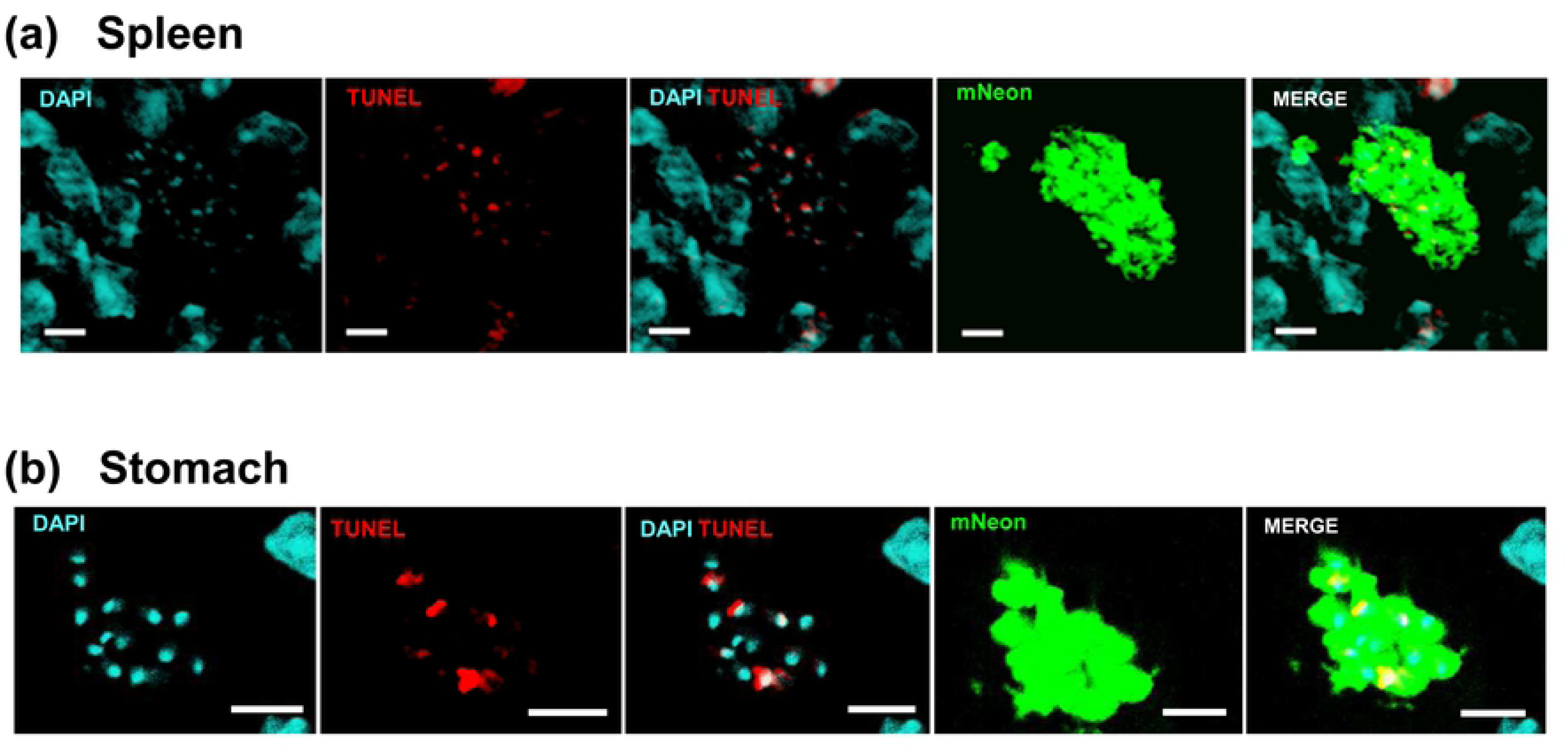
Asynchronous replication of parasite mitochondrial DNA within single infected host cells *in vivo* revealed by TUNEL assays. (a) Asynchronous replication of kDNA in intracellular parasites infecting mouse spleen cells during an acute stage infection (day 19). BALB/c mice were infected with *T. cruzi* CL-Luc::Neon and histological sections prepared from bioluminescent tissue (Experimental procedures). Parasites were detected by green fluorescence (mNeon), and the tissue sections subjected to TUNEL assays to highlight replicating kDNA (red). (b) Asynchronous replication of kDNA in an amastigote nest detected in the smooth muscle layer of stomach tissue during a chronic stage infection (day 117). Bar = 10 μm.

**FIG 3.**
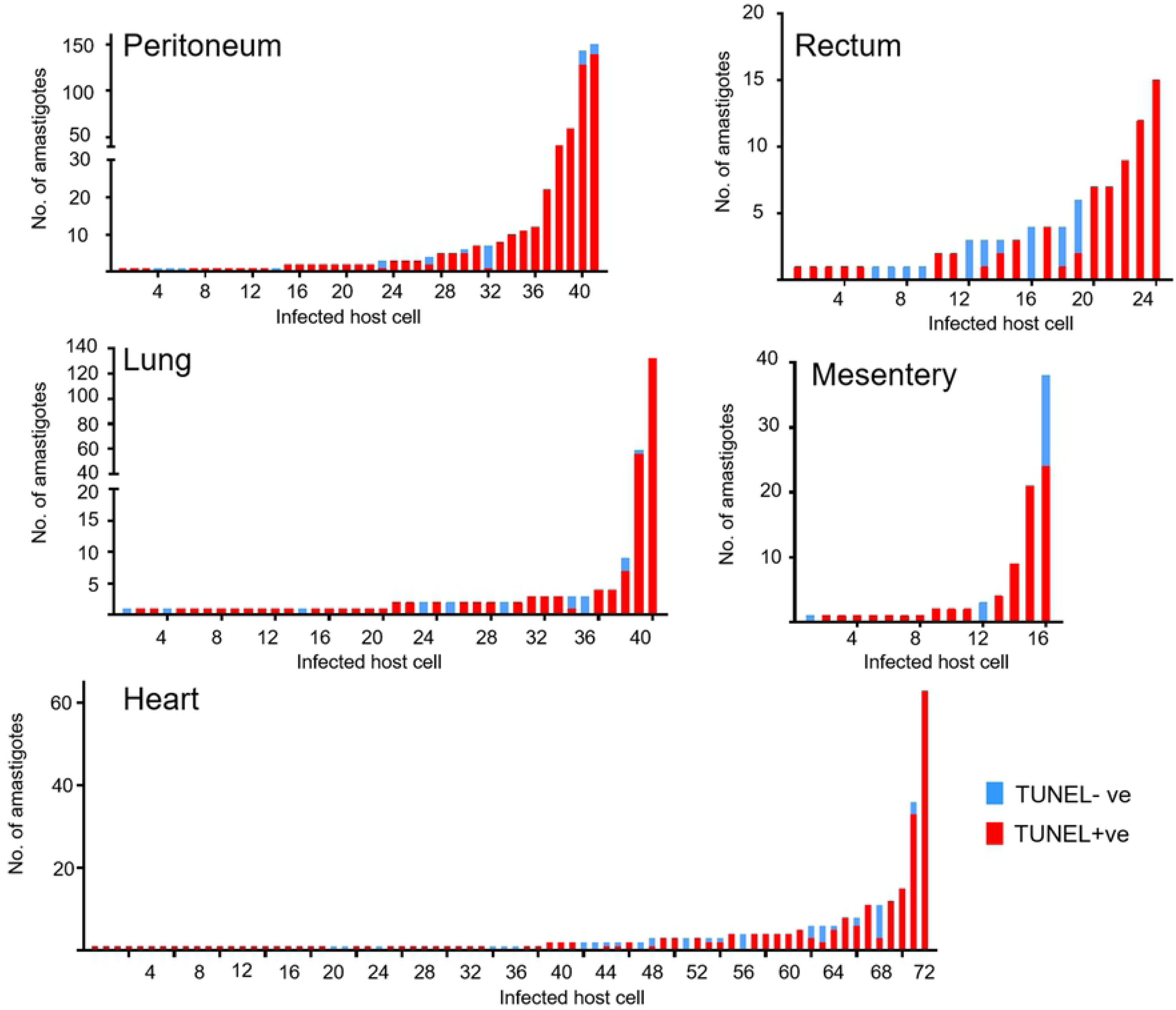
Quantification of TUNEL in BALB/c mice during the acute stage of infection with *T. cruzi* CL-Luc::Neon. Tissue sections from mice sacrificed on day 19 post-infection were processed for imaging and subjected to TUNEL staining (Experimental procedures). The graphs show the number of amastigotes that were TUNEL+ve (red) or TUNEL-ve (blue) in individual infected cells within the specified tissues. The x-axis refers to individual host cells. Bars containing both TUNEL-ve and +ve amastigotes were present in all tissues examined. Note that the level of TUNEL signal may vary between amastigotes within a given cell, so even bars that are red only may represent parasites at different stages of kDNA replication (c.f. differential levels of TUNEL staining in Fig 2a and b, DAPI/TUNEL panels).

### Replication of parasite nuclear DNA is not synchronised within individual infected host cells

TUNEL assays identify parasites where kDNA replication has initiated, but do not provide information on those where it has terminated and the parasite has progressed to nuclear DNA synthesis. To get a more quantitative picture of both nuclear and kinetoplast replication, we injected *T. cruzi*-infected mice with the nucleoside analogue 5-ethynyl-2’-deoxyuridine (EdU) at specific time points prior to necropsy [29]. We chose EdU rather than BrdU, since this analogue can be fluorescently labelled directly in double stranded DNA and does not require harsh denaturing conditions. This preserves the mNeonGreen fluorescence used to locate *T. cruzi in situ*. EdU is incorporated into newly synthesised DNA molecules and identifies parasites undergoing nuclear or kDNA replication during the time period of the EdU pulse. It also labels mammalian cells that enter S-phase during this period. EdU distribution in murine tissues is extensive and incorporation is stable. For example, Merkel cells from mice whose mothers were injected with EdU during pregnancy remain labelled nine months after birth, suggesting that the analogue is not removed during DNA repair [30-32]. Labelling of replicating host cells within a given tissue section can therefore be used as an internal control for EdU tissue penetration to sites of *T. cruzi* infection. Fixed tissue sections containing host cells and/or parasites that incorporate EdU are fluorescently labelled by click chemistry and can be examined by confocal microscopy [33] (Experimental procedures).

We assessed a range of bioluminescence positive tissues excised from mice in the acute stage of infection (Fig 4). In cardiac sections, there was negligible labelling of host cell nuclei, as expected, since heart muscle consists predominantly of terminally differentiated non-replicative cells. However, labelled intracellular parasites were easily detected. Within host cells containing multiple parasites, EdU labelling was heterogeneous across the population and many parasites had not incorporated EdU (Fig 4a) during the time of exposure. Similarly, in adipose tissue, parasites within the same infected cells displayed a wide range of EdU specific fluorescence intensity (Fig 4b). This heterogeneity was dispersed throughout the infected cell, with replicating and non-replicating organisms being interspersed.

**FIG 4.**
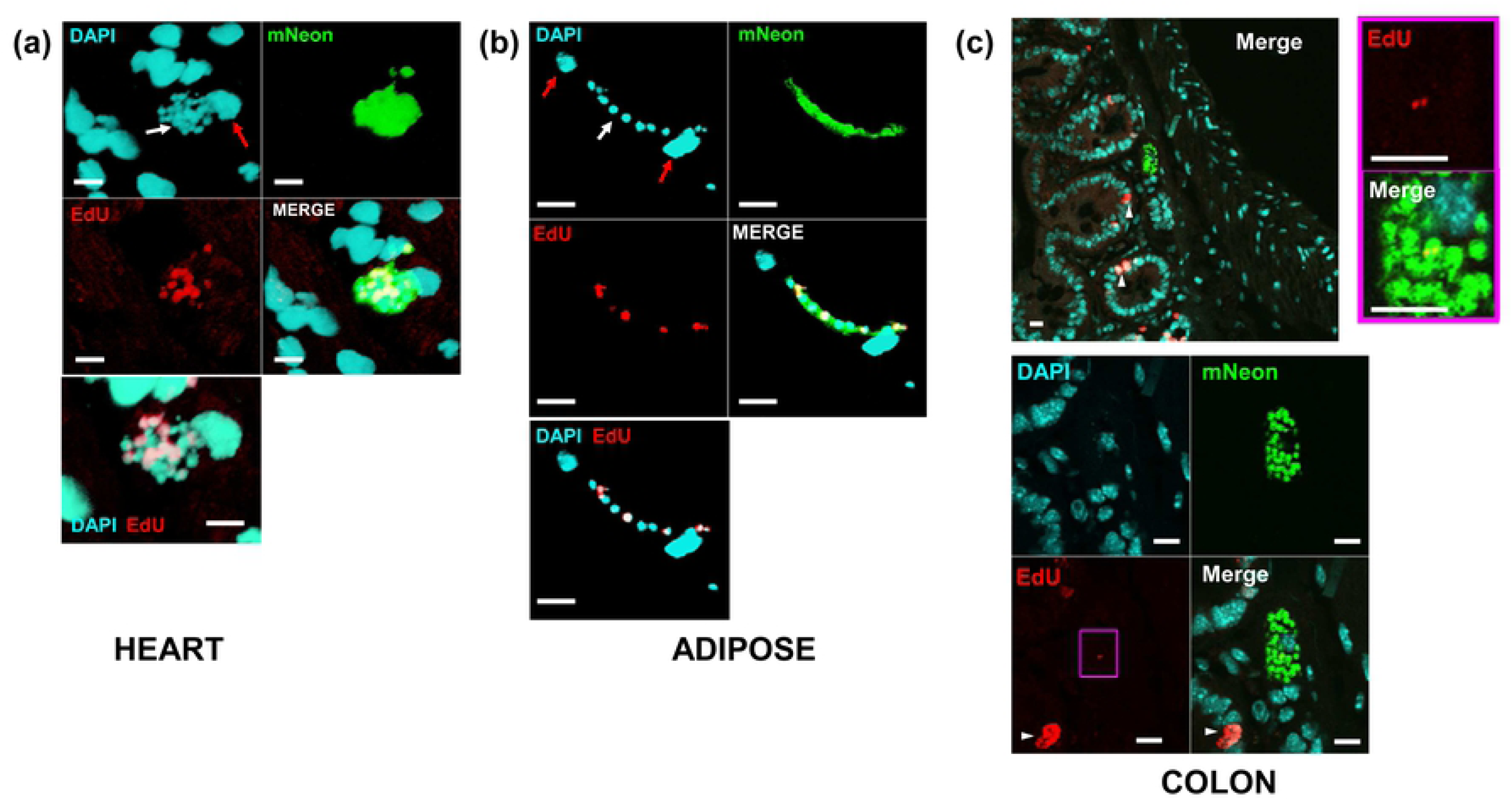
Asynchronous parasite DNA replication within single infected host cells *in vivo* revealed by EdU-labelling. Replication of parasite DNA within mice infected by *T. cruzi* clone CL-Luc::Neon was assessed after inoculating EdU (for (a) and (b), one pulse 6 hours prior to tissue sampling; for (c), two pulses 18 and 28 hours prior to tissue sampling (Experimental procedures). Parasite location in histological sections was detected by green fluorescence (mNeon). (a) DNA replication (EdU, red) in a parasite nest during an acute stage infection (heart tissue, day 15 post-infection). In the DAPI stained image, the white arrow indicates parasite nest, and red arrow the host cell nucleus. The merged DAPI/EdU image, bottom left, illustrates the heterogeneity in the DNA replication status of parasites within the nest. (b) DNA replication in parasites within adipose tissue (day 15 post-infection). Red and white arrows in the DAPI image identify host and parasite DNA, respectively. Combined EdU and DAPI image shows replicating parasites interspersed with non-replicating parasites. (c) Section from GI tract of mouse, upper panel shows image at low magnification – note the presence of some EdU+ve mammalian cells within the mucosal layer due to epithelial cell replacement (indicated by white arrowheads). Lower panels show magnified view of parasite nest. EdU signal in magenta box is shown in higher magnification to the right; note a single amastigote with EdU labelling at antipodal sites of kDNA replication. All other parasites in this nest are negative. Bars = 10 μm.

In gut sections obtained from chronically infected mice, EdU labelling of host cells in the mucosal epithelium was readily apparent, since these cells are continually shed into the gut lumen and replaced from stem cells (Fig 4c, white arrowheads). As in the acute stage, the labelling pattern within amastigote “nests” was consistent with asynchronous replication of nuclear DNA, with many parasites showing no detectable EdU incorporation (Fig 4c; S4 Fig a, b). We also analysed sections taken from tissue samples that contained all of the detectable bioluminescent foci in the gastrointestinal tract of three individual chronically infected C3H/HeN mice (M275-17, M277-17 and M279-17). We injected these animals with two pulses of EdU at 18 and 28 hours before euthanasia. The number of parasites and infected cells was consistent with the strength of the bioluminescent signal visible on *ex-vivo* organ sections (Fig 5a). Some of the nests were very large (“mega-nests”), containing hundreds of parasites, and in some cases, they clearly extended beyond the limits of the tissue section (indicated by asterisks, Fig 5b, c). However, examination of serial sections of a single large nest indicated that the asynchronous nature of EdU incorporation was sustained throughout the nest (Fig 6), since in each section there were both EdU+ve and Edu-ve amastigotes. The extent of EdU labelling within amastigotes in an infected cell was variable as had been observed with the TUNEL assay. This would be expected if parasites were sampled at different stages within S-phase. It was clear that many parasites had not replicated during the period of EdU exposure because most amastigotes (77% in the GI tract, 62% in the peritoneal muscle) were negative for EdU labelling in either kinetoplast or nucleus. Therefore, both TUNEL assays and EdU incorporation demonstrate that *in vivo*, the timing of DNA replication is autonomous to individual parasites within an infected host cell, with no evident synchronisation of the process between different amastigotes.

**FIG 5.**
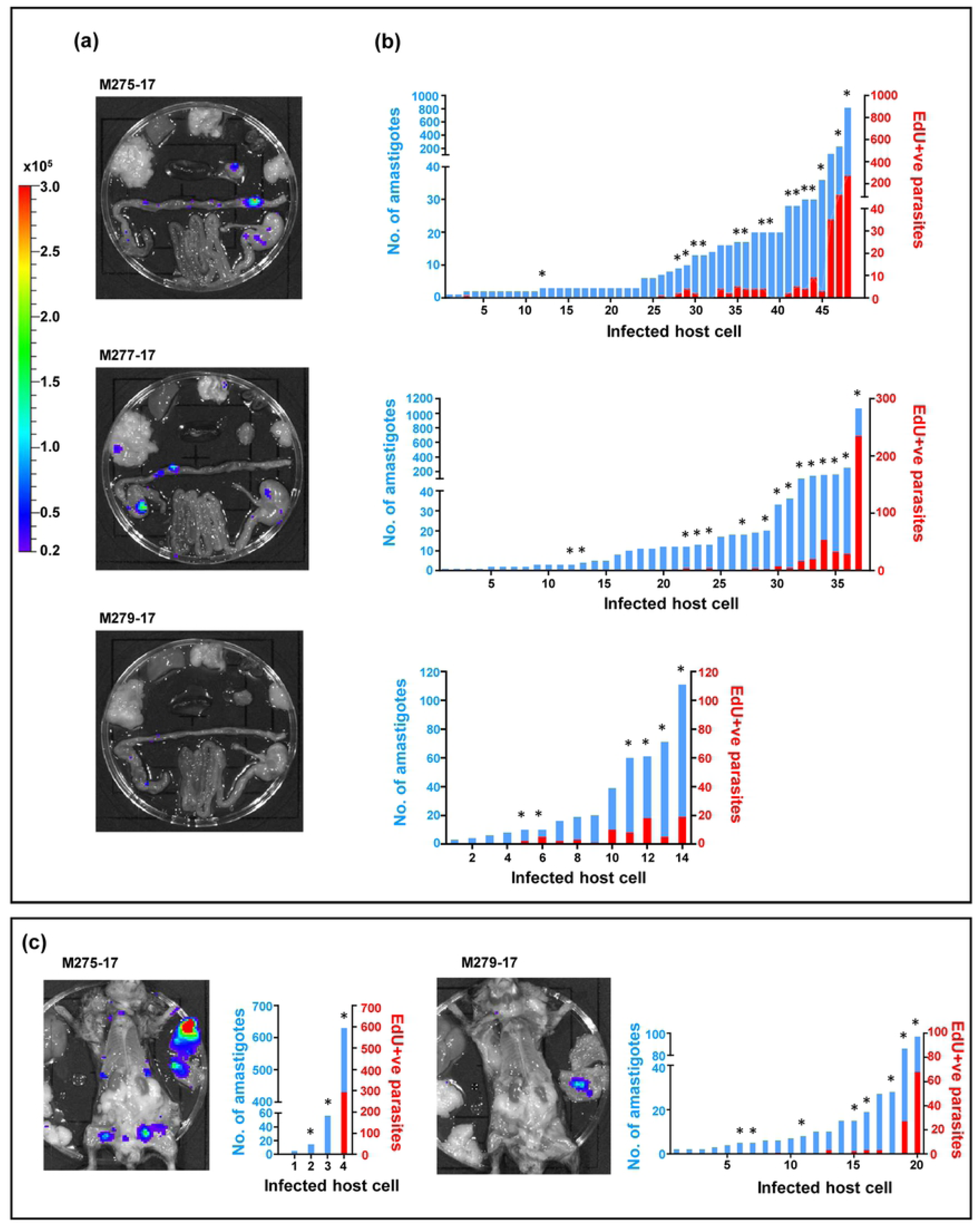
EdU labelling reveals that cells infected with small numbers of amastigotes have a lower percentage of actively replicating parasites in a chronic infection. (a) *Ex vivo* imaging of organs. Bioluminescent foci were removed from the GI tract of three chronically infected C3H/HeN mice (day 211 post-infection) that had been injected with two pulses of EdU 18 and 28 hours prior to necropsy (Experimental procedures). (b) Each infected cell in the GI tract foci was imaged and the number of amastigotes that were positive or negative for EdU incorporation was quantified. The graphs show the total number of amastigotes in each cell (blue bars) and the number that were labelled with EdU (red bars). (c) Bioluminescent foci from the peritoneal muscle were also dissected, stained for EdU and quantified as above. Asterisks above bars indicate cells were the number of parasites represents a minimum due to the infected nest being larger than the z-dimension of the section.

**FIG 6.**
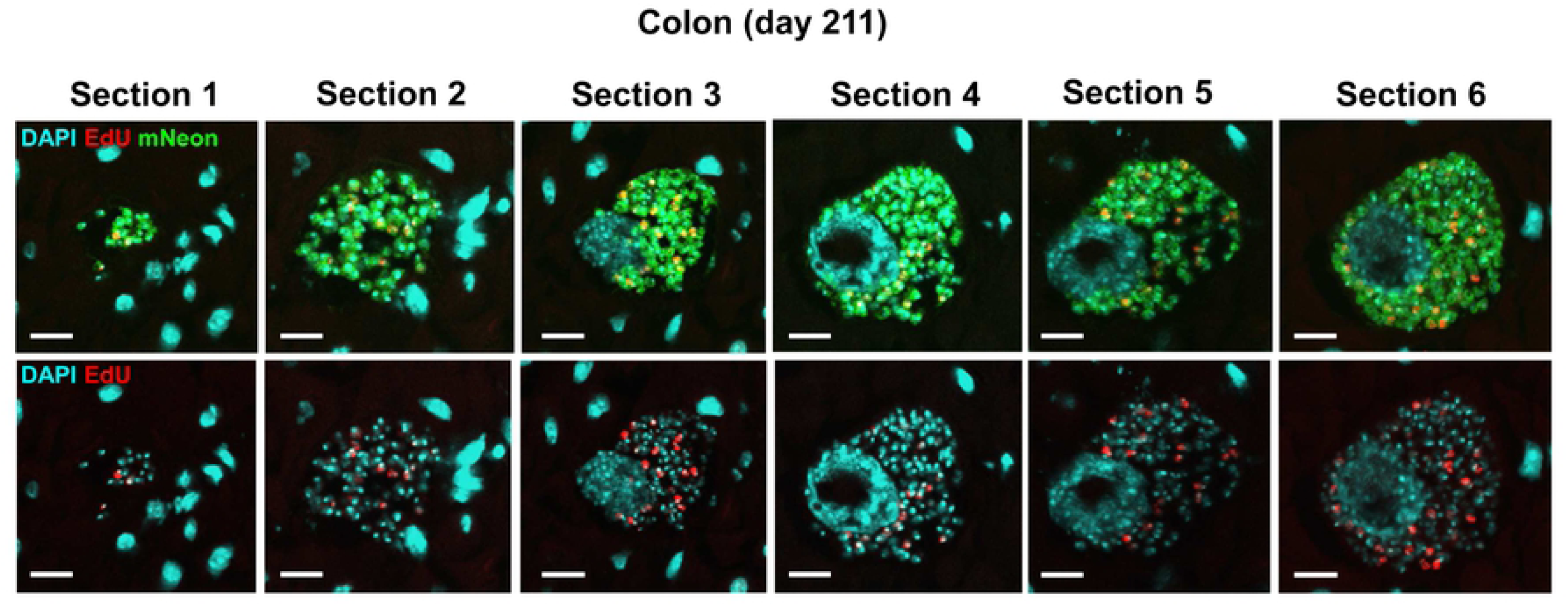
Large nests are present in the chronic stage of infection (C3H/HeN mouse, day 211) and show asynchronous EdU incorporation throughout. Images of the same nest taken from different sections through the tissue. The top row shows DAPI, EdU and mNeonGreen merged channels, whilst the lower row shows DAPI and EdU channels (for clarity). Bar = 10 μm. Note that sections are from the same infection focus but not all sections of this nest are included due to loss in processing.

### Both replicating and differentiating parasites co-exist in the same host cell

The final step in the intracellular development of *T. cruzi* is differentiation of replicating amastigotes into non-dividing flagellated trypomastigotes, prior to their escape from the host cell. The mechanisms that regulate this process *in vivo*, from a temporal and organisational perspective, are unknown. In mammalian cell monolayers infected *in vitro*, we observed that amastigotes could be detected in the same cells as differentiated trypomastigotes (Fig 7a). We used the TUNEL assay to examine whether amastigotes in this environment were undergoing replication or were about to differentiate. Antipodal TUNEL staining was observed in the kinetoplasts of some amastigotes present in cells with trypomastigotes indicating ongoing kDNA replication (Fig 7b). Co-existence of replicating parasites with trypomastigotes was confirmed by live-cell imaging of infected cells *in vitro* (S5 Fig, S1 Movie, S2 Movie). This suggested asynchronicity in the process of both differentiation and cell division. Amastigotes can therefore initiate a new replicative phase while in the same host cell as parasites that have differentiated to trypomastigotes as judged by morphology and flagellar position. It remains possible that some amastigotes initiate replication but then “pause”, leading to TUNEL+ve parasites co-existing with flagellated trypomastigotes.

**FIG 7.**
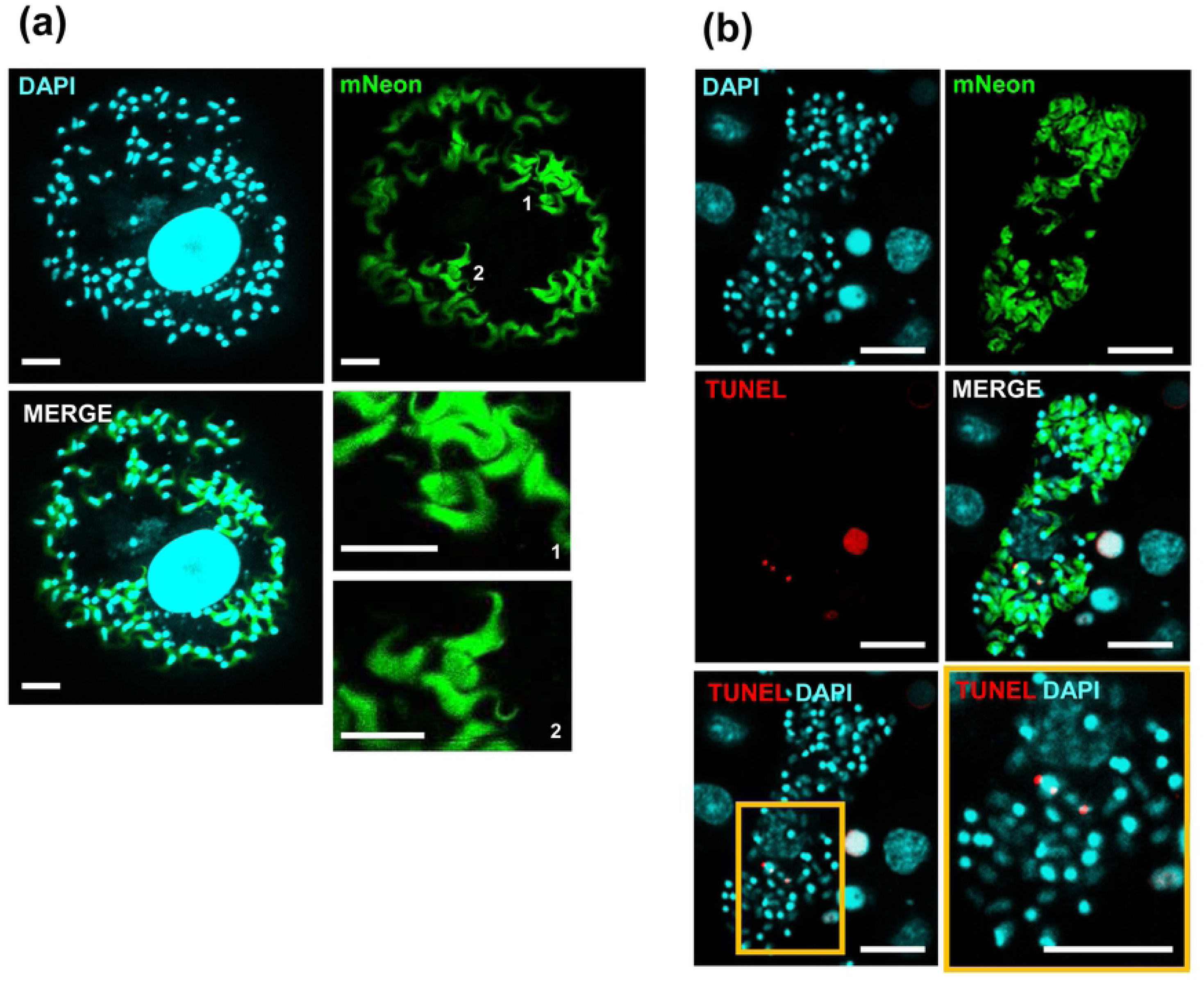
TUNEL assays indicate that amastigote replication and amastigote-to-trypomastigote differentiation can occur concurrently within single infected host cells *in vitro*. (a) MA104 cells infected *in vitro* with *T. cruzi*. Two amastigotes (1 and 2) are visible within a cell full of trypomastigotes. The two lower right-hand panels show the two amastigotes at a higher magnification for clarity. (b) MA104 cells infected *in vitro* with *T. cruzi*. The cells were fixed 72 hours post-infection and subjected to a TUNEL assay. Two replicating amastigotes can be identified by antipodal TUNEL labelling on the kinetoplast, amongst a population of differentiated trypomastigotes. Bar = 10 μm.

### Multiple morphological forms of *T. cruzi* are present in deep tissues of infected mice

Classically, the *T. cruzi* life-cycle in mammals involves two distinct morphological stages, the intracellular replicative amastigote, which lacks an external flagellum, and the non-replicating extracellular flagellated trypomastigote. However, other forms of the parasite have been observed under *in vitro* conditions (for review, [34]). These observations normally involve only one host cell type, and lack environmental signals and a tissue milieu. Therefore, it has not been possible to be assess if these non-classical forms are physiologically relevant during host infections, or whether they are artefacts of *in vitro* culture.

We observed a number of distinct *T. cruzi* morphological forms during murine infections that do not conform to the standard amastigote/trypomastigote dichotomy. In both acute and chronic infections, we frequently visualised amastigote-like forms with a protruding flagellum (Fig 8). This flagellum extended from the anterior of the parasite, based on the relative position of the kinetoplast and nucleus (Fig 8a-c). The kinetoplast and nucleus displayed the forms associated with the replicative stages of the parasite. The length of the visible flagellum was highly variable with the majority of amastigotes having no protruding flagellum. (Fig 8d). The length of the amastigote cell body varied between 3 and 7 µm (mean 4.2 ± 0.8 µm) with the flagellar length being independent of cell body length (Fig 8c and e). The flagellated amastigote-like parasites have similarities to sphaeromastigotes (Tyler & Engman, 2001), a form that has been observed *in vitro*.

**Fig 8.**
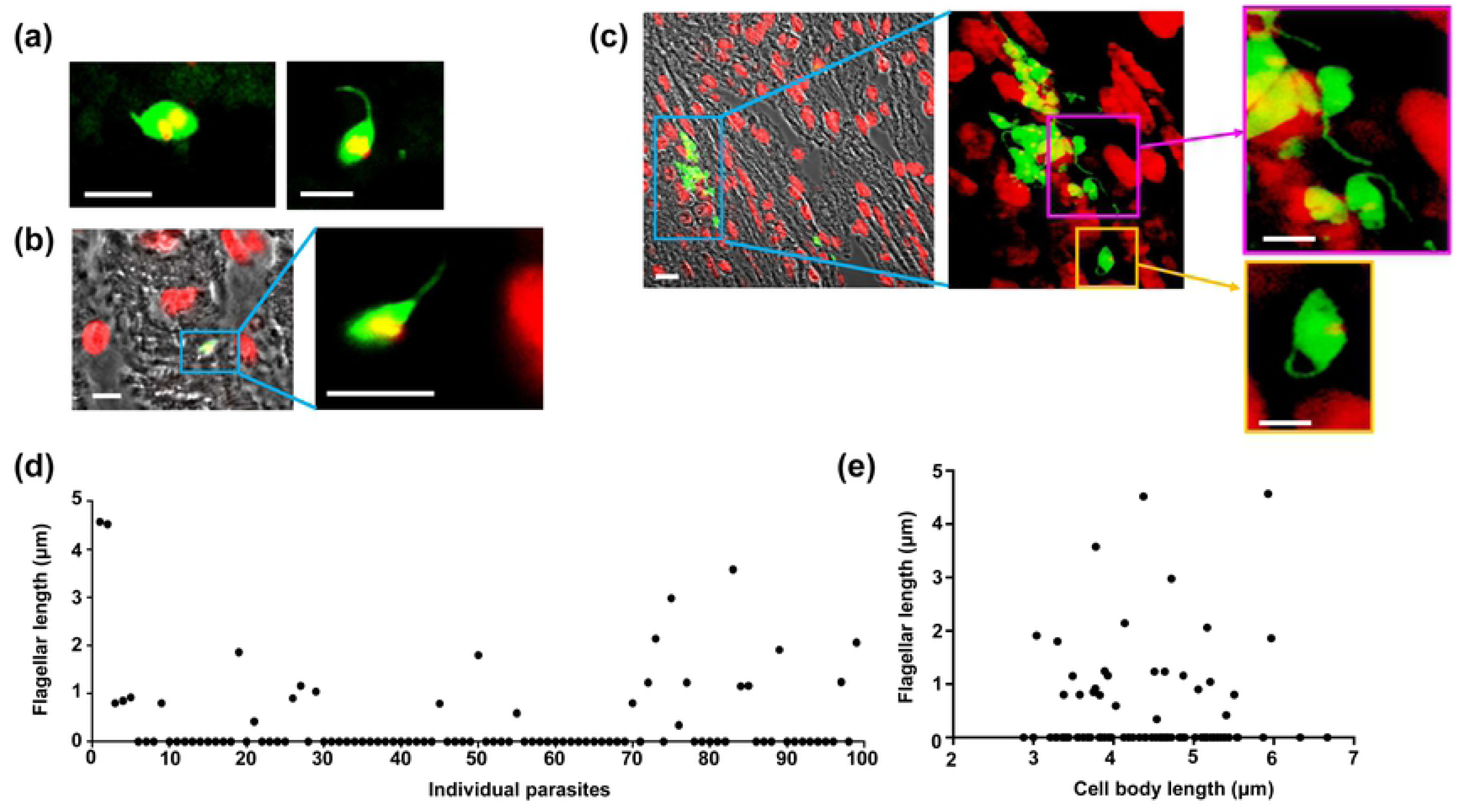
*T. cruzi* parasites display a wide range of morphologies during murine infections. BALB/c mice were inoculated with parasites expressing a fluorescent/bioluminescent fusion protein and infected tissues identified by *in vivo* bioluminescence imaging (Experimental procedures). Fluorescent (green) flagellated “amastigote” forms detected in (a) adipose tissue (day 13 post-infection) (DNA stained red – appears yellow where mNeon fluorescence overlaps DNA), and (b) cardiac tissue (day 19 post-infection). (c) Parasite nests in the rectum (day 19 post-infection) containing a variety of morphological forms. Note that none of the flagellated forms displays the posterior rounded kinetoplast characteristic of trypomastigotes. Bar = 5 μm. (d and e) The flagellar length was measured in 100 amastigote-like cells from various tissue sites, where parasites were distinct enough to measure both flagellum and cell body. (d) Graph showing the flagellar length (μm) measured in each individual amastigote. (e) Graph showing the flagellar length (μm) plotted against parasite body length (μm).

In addition to the flagellated amastigote-like parasites, we also observed a second non-standard form that displays an epimastigote-like morphology (Fig 8c, orange box and inset). Similar forms have been reported once before in a very early stage of infection (day 8) [35]. These epimastigote-like forms, which we detected repeatedly in tissue samples, often co-existed with dividing amastigotes and differentiating trypomastigotes in the same infected cell, and could be observed by live cell imaging *in vitro* (S5 Fig). Whether these forms are simply morphological intermediates, or have a distinct role in infection or transmission remains unknown.

## DISCUSSION

The broad outline of *T. cruzi* replication and stage-specific differentiation during mammalian infection has been known for more than a century. However, it is clear that this part of the life-cycle is more complex than previously described, with possible implications for our understanding of pathogenesis, immune evasion and transmission [34]. Unravelling the biology of *T. cruzi* within the host is also crucial from a drug development perspective, since some life-cycle stages may be less sensitive to treatment [9], and the ability of the parasite to reside in metabolically distinct tissue compartments may have significant effects on drug exposure and pharmacodynamics. To date, most research on *T. cruzi* replication and differentiation has utilised *in vitro* systems. Although these are informative, they may not capture the full developmental range, and could give rise to artefactual observations that are not relevant to these processes within the mammalian host. In addition, *in vitro* cultures often use immortalised mammalian cell lines, whereas *in vivo T. cruzi* is usually found in non-replicating terminally differentiated cells such as muscle fibres.

One of the major unknowns in *T. cruzi* biology is the extent to which parasite growth is co-ordinated within individual host cells during a mammalian infection, and how it is influenced by tissue/organ location and disease status. This issue has been highlighted by recent reports of spontaneous dormancy during intracellular infection (Sánchez-Valdéz et al., 2018). Here, using a bioluminescent/fluorescent dual reporter strain that significantly enhances our ability to identify and visualise infected host cells *in vivo*, we provide evidence that intracellular replication is largely asynchronous. From observation, it is apparent that the number of parasites per host cell does not follow a predictable or tightly regulated pattern *in vitro* (Fig 1, S2 Fig), or *in vivo*, at any phase of the infection, or in any specific tissues (Figs 2-6). Consistent with this, two separate assays indicate that, within individual infected cells, DNA replication is not synchronised between parasites at either nuclear or kinetoplast genome levels (Figs 2-6, S3, S4). In the case of EdU labelling, this was not a reflection of differential tissue penetration, since replicating amastigotes were interspersed with non-labelled parasites in a wide range of tissues types, during both acute and chronic infections. TUNEL labelling is not dependent on incorporation of nucleoside analogues in a living mouse and is therefore an orthogonal assay for mitochondrial DNA replication.

The finding that extremely large nests of asynchronously dividing or differentiating parasites can exist in chronically infected animals (Fig 6 and S4 Fig) could have therapeutic implications. Infected cells such as these may contain parasites in a range of metabolic states (including dormancy) that exhibit heterogeneity in terms of drug susceptibility. In addition, the possibility that these *in vivo* mega-nests could result in some form of intracellular “herd-protection” may give rise to an environment that is difficult to replicate in the standard *in vitro* assays used in the drug development pipeline.

Single infected cells can contain both replicating amastigotes and non-replicating, differentiated trypomastigotes (Fig 7). Therefore, whatever the signal(s) that trigger differentiation and/or replication, they are not perceived and/or acted on in concert by every parasite within the nest. This contrasts with the related extracellular parasite *T. brucei* in which a well-characterised quorum sensing pathway initiates differentiation from the replicative long slender bloodstream form to the non-replicating short stumpy form, preadapted for transmission to the tsetse fly vector [36-39]. The lack of synchrony in differentiation between amastigote, intracellular “epimastigote” and trypomastigote, during *T. cruzi* infection, indicates that either a ubiquitous quorum sensing mechanism of this kind does not operate within single infected host cells, or that some parasites remain refractory to the trigger signal, as exemplified by the quiescent amastigotes identified recently [9].

The dual reporter parasite strain also enabled us to identify a number of non-standard parasite forms in tissues of infected mice, sometimes co-existing within the same host cell (Fig 5, S5 Fig). The role of the intracellular and extracellular epimastigote-like, and flagellated amastigote-like forms in the parasite life-cycle remains to be determined. Their relative scarcity suggests that they could be transient forms which occur during the differentiation from amastigote to trypomastigote. Importantly, detection of these morphological forms *in vivo* excludes the possibility that they represent laboratory culture artefacts. Intriguingly, in this context, it has been established that in the opossum, an ancient natural host of *T. cruzi*, there is an insect stage-like epimastigote cycle within the anal glands. This appears to exist independently of the intracellular pathogenic cycle found in other tissues [40]. It has also been demonstrated that trypomastigotes can exist in two distinct populations (TS+ and TS-, referring to *trans*-sialidase surface expression). TS-parasites are poorly infective to mammalian cells and significantly less virulent in mice [41]. This suggests that the two populations may have distinct roles, one perhaps preadapted for invasion of the insect vector, and the other for propagation of infection within the mammalian host, analogous to the slender and stumpy forms of *T. brucei.*

In conclusion, this study reports the first detailed analysis of *T. cruzi* replication in animals at the level of single infected cells within a range of tissue types. The data reveal the complexity of parasite replication and differentiation cycles, and confirm the existence *in vivo* of parasites with a non-classical morphology. The presence of even transient non-canonical forms in infected animals highlights important questions about their susceptibility to trypanocidal drugs, compared with standard amastigotes. Similarly, it is unknown whether these forms express the same surface protein repertoire as amastigotes and/or trypomastigotes, if they are equally targeted by anti-parasite antibodies in the bloodstream and tissue fluids, or if they retain the ability to infect other cells and disseminate the infection. It will now be important to develop procedures to isolate these non-classical parasite types in sufficient numbers to allow their biochemical and biological characterisation.

---

**S1 Fig** MA104 cells infected with *T. cruzi* CL-Luc::Neon amastigotes for 72 hours, fixed, then labelled with the TUNEL reagent. The parasite in the red box has completed kDNA replication and segregation, but not nuclear replication, and clearly shows that the segregated kinetoplasts no longer display TUNEL positivity.

**S2 Fig** Plot of TUNEL+ve amastigote numbers as a function of total amastigotes present in an infected cell, for each infected cell used to derive Fig 1d. Each circle represents a single infected host cell. (a) All 200 infected cells from Fig 1d. (b) An expanded view of the area indicated by the box to allow clear visualisation of the host cell numbers. For cells infected with 1 amastigote, n=28.

**S3 Fig** Asynchronous parasite kDNA replication within single infected host cells *in vivo* in acutely infected (19 days post infection) BALB/c mice revealed by TUNEL reactivity. (a) caecum, (b) rectum, (c) heart, (d) spleen and (e) lung. Images are from two individual mice. Bar = 10 μm

**S4 Fig** Asynchronous parasite DNA replication within single infected host cells in chronically infected (211 days post infection) C3H/HeN mice revealed by EdU-labelling. Replication of parasite DNA within mice infected by *T. cruzi* clone CL-Luc::Neon (Costa et al., 2018) was assessed after inoculating two EdU pulses 18 and 28 hours prior to tissue sampling (Experimental procedures). Parasites were located in histological sections by fluorescence (mNeon, green). a) DNA replication (red) in a chronic phase parasite nest (colon). The combined DAPI/EdU image illustrates the heterogeneity of parasite replication within the nest. Bar = 10 μm. b) Section from colon of mouse showing parasite nest. Upper panels show individual channels and a merged image. The lower panel shows DAPI and EdU channels only, allowing visualisation of the interspersed nature of EdU+ve amongst EdU-ve parasites. (a) and are from different mice. Bars indicate 10 μm.

**S5 Fig** Multiple morphological forms within single infected cells. Each image shows an M104 cell (blue, nucleus) 6 days after infection with *T. cruzi* (green) showing amastigotes (arrow a) dividing amastigotes (arrow da), epimastigote-like forms (arrow e) and trypomastigotes (arrow t) within the same cell. (a-d) sequential still images from S1 Movie, (e-h) sequential still images from S2 Movie. Bars indicate 20 µm.

**S1 Movie** Multiple morphological forms within a single infected cell Live cell imaging of an M104 cell 6 days after infection with *T. cruzi* showing dividing amastigotes, epimastigote-like forms and trypomastigotes within the same cell. See S5 Fig a-d for locations of representative parasites for each morphotype.

**S2 Movie** Multiple morphological forms within a single infected cell. Live cell imaging of an M104 cell 6 days after infection with *T. cruzi* showing amastigotes, epimastigote-like forms and trypomastigotes within the same cell. See S5 Fig e-h for locations of representative parasites for each morphotype.

